# Links between environment, diet, and the hunter-gatherer microbiome

**DOI:** 10.1101/319673

**Authors:** Gabriela K. Fragiadakis, Samuel A. Smits, Erica D. Sonnenburg, William Van Treuren, Gregor Reid, Rob Knight, Alphaxard Manjurano, John Changalucha, Maria Gloria Dominguez-Bello, Jeff Leach, Justin L. Sonnenburg

## Abstract

The study of traditional populations provides a view of human-associated microbes unperturbed by industrialization, as well as a window into the microbiota that co-evolved with humans. Here we discuss our recent work characterizing the microbiota from the Hadza hunter-gatherers of Tanzania. We found seasonal shifts in bacterial taxa, diversity, and carbohydrate utilization by the microbiota. When compared to the microbiota composition from other populations around the world, the Hadza microbiota shares bacterial families with other traditional societies that are rare or absent from microbiotas of industrialized nations. We present additional observations from the Hadza microbiota and their lifestyle and environment, including microbes detected on hands, water, and animal sources, how the microbiota varies with sex and age, and the shortterm effects of introducing agricultural products into the diet. In the context of our previously published findings and of these additional observations, we discuss a path forward for future work.

## The seasonality of the Hadza microbiota

We recently characterized the gut microbiota of the Hadza hunter-gatherer population(*1*). The Hadza live in the Central Rift Valley in Tanzania and have historically subsisted on five groups of foraged and hunted foods: berries, honey, baobab, tubers, and meat(*2*). The Hadza experience two main seasons: wet (November to April) and dry (May to October). These seasons are accompanied by shifts in available food and activities. For example, while hunting occurs throughout the year, meat is taken more often in the dry season when water sources are more predictable and ambush-versus encounter-hunting can be practiced more frequently. Conversely, more honey is eaten during the wet season. Fiber-rich tubers are eaten throughout the year.

We found that the composition of sampled gut microbial communities from the Hadza corresponded with seasonality. Fecal samples were taken during the dry season in 2013, the following wet season in 2014, and the following dry season in 2014. Of 350 fecal samples, 188 were used in the analysis, each from a different individual to avoid bias from repeated sampling of the same individual. The composition of the wet season community was distinct from that of both dry seasons, whereas the dry season-compositions of 2013 and 2014 were statistically indistinguishable from one another. Samples taken from a previous study of the Hadza microbiota during the early wet season in 2013(*3*) were consistent with this pattern.

To understand this cyclic pattern, we tracked operational taxonomic units (OTUs) across the seasons. We found that OTUs from the phylum Firmicutes exhibited relative stability across seasons compared to the Bacteroidetes, a phylum in which half of OTUs were lost during the wet season and reappeared during the following dry season. Both in the full dataset of 188 individuals, and in a subset of individuals sampled longitudinally across the three studied seasons, shifts in OTU abundance occurred primarily in four bacterial families: Prevotellaceae, Succinovibrionaceae, Paraprevotellaceae, and Spirochaetaceae. The seasonal pattern of one family, the Prevotellaceae, is shown in the cross-sectional population in Figure 1A.

**Figure 1:**
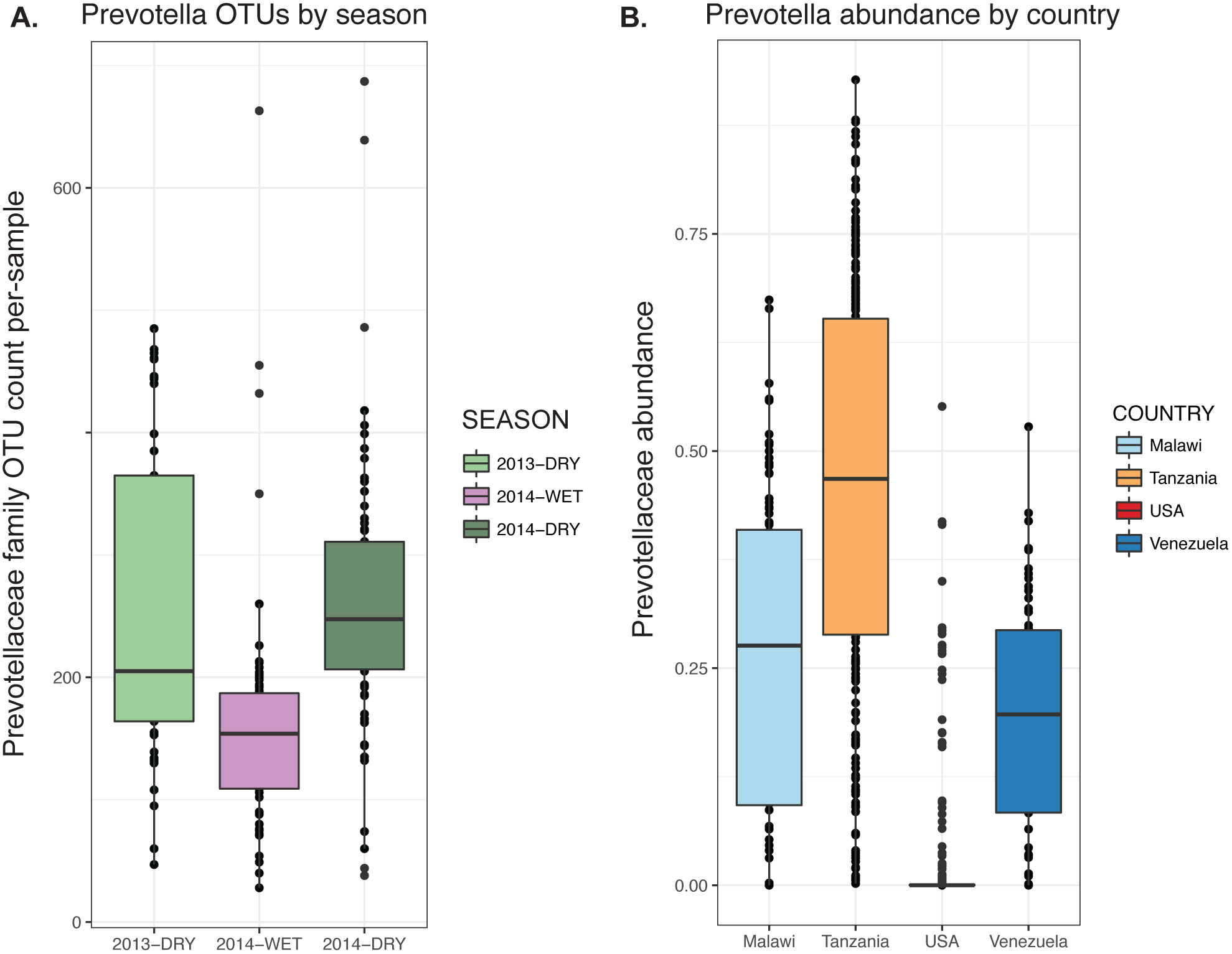
Seasonally volatile bacterial families in the Hadza are prevalent in traditional populations and diminished in the industrialized microbiota. A. The number of OTUs from the Prevotellaceae family observed per sample in 188 Hadza fecal samples, partitioned by season (2013-Dry, n = 41; 2014-Wet, n = 77; 2014-Dry, n = 70). Data rarefied to 11,000 OTUs.
B. Relative abundance of Prevotellaceae family in samples populations in rural Malawi, Tanzania (Hadza hunter-gatherers), metropolitan area in USA, and Amazonas in Venezuela.

We also discovered differences in carbohydrate utilization capacity of the Hadza gut microbiota across seasons. The analysis of genes encoding carbohydrate-active enzymes (CAZymes)(*4*) present in metagenomic data from the same Hadza individuals sampled across three seasons revealed a cyclical pattern of CAZyme diversity, with higher diversity observed during the dry season. There were no differences between the dry seasons in consecutive years, but samples from the wet season had lower levels of CAZymes that degrade carbohydrates from animal, plant, and mucin sources. These seasonal shifts in the microbiota’s capacity to process different types of carbohydrates may reflect seasonal dietary differences. While broad seasonal dietary trends have been documented among the Hadza(*2*), a more thorough and nuanced analysis of how specific dietary patterns connect to gut microbiota dynamics is needed.

Comparisons between the Hadza and an American cohort from the Human Microbiome Project(*5*) revealed significant differences. The Hadza gut microbes possess higher levels of genes encoding plant-degrading enzymes whereas genes encoding enzymes targeted to animal and mucin degradation were enriched in the American cohort. This distinction is consistent with differences in diet between Americans and Hadza. The Hadza diet is rich in microbiota-accessible carbohydrates (MACs) found in plant-based dietary fiber, however the MAC-poor American diet(*6, 7*) selects for gut microbes well adapted to forage on intestinal mucus(*8–10*).

## A microbiota conserved across traditional populations is lost in industrialized nations

We combined the Hadza microbiota data with data from 18 different populations from 16 countries, comprising a variety of lifestyles including hunter-gatherer, agrarian, and industrialized(*11–16*). When combined into a Bray-Curtis dissimilarity PCoA plot, reflecting the degree of shared taxa between samples, microbial composition data from industrialized cohorts separated from the traditional cohorts, the first principal component capturing a gradient of modernization. An examination of the bacterial families that differed in abundance across these populations revealed that traditional populations tended to have higher levels of Prevotellaceae, Succinovibrionaceae, Paraprevotellaceae, and Spirochaetaceae, whereas industrialized cohorts had higher levels of mucus-consuming Verrucomicrobia and Bacteroidaceae. As an example, we show in the abundance of Prevotellaceae across four countries (Figure 1B), which reveals variation that is striking for two reasons. Firstly, while differences between the microbiotas from industrialized and traditional populations may have been expected, the conservation of higher levels of Prevotellaceae and other taxa across geographically separated traditional populations suggests that these organisms have evolved as particularly well-adapted to its human host. Rather than being a feature of a specific geographical environment, the global pervasiveness and association of these microbes with a lifestyle that defined humans for much of our existence as a species suggests the functions associated with these taxa may have shaped human biology, and appear to have been lost through industrialization. Secondly, the bacterial families that differentiate traditional and industrialized populations also exhibit seasonal volatility in the Hadza. This finding indicates that volatility in abundance may serve as a marker of microbes that are vulnerable to eradication via various perturbations including those associated with modernization. These factors, such as increased sanitation and hygiene, may further act as barriers to reacquiring these lost microbes, such that these losses persist. The role of the families that have been lost or have become rare within the industrialized intestinal ecosystem and their interactions with the host is an open and important question. In the remaining text, we will use the term VANISH (Volatile, Associated Negatively with Industrialized Societies of Humans) to refer to taxa in the families Prevotellaceae, Succinovibrionaceae, Paraprevotellaceae, and Spirochaetaceae.

## The Hadza and their environment: microbes associated with hands, animals, and water sources

Environmental exposure to microbes varies across lifestyles and certainly between traditional and industrialized populations. We were curious whether aspects of the Hadza lifestyle, particularly interaction with the natural environment through hunting, foraging, and drinking untreated, surface water sources may serve as a reservoir of bacterial diversity for the microbiota. We were fortunate to have access to additional samples that we assessed for bacterial composition, and we focused our analysis on the gut-associated bacterial families identified as seasonally volatile and conserved in traditional societies.

The skin on the right hand of individuals were sampled using swabs during the dry season. A PCoA plot of weighted Unifrac distance of hand and fecal samples shows a separation by body site, consistent with observations in industrialized populations that also distinguish samples by body site (Figure 2A)(*17, 18*). In addition, the hand samples (dry season), are clustered closer to the fecal samples taken from the dry season than fecal samples from the wet season. Although hand samples from the wet season were not available as a point of comparison, the data suggest a degree of concordance between the hand and fecal samples by season. Examining the VANISH microbial families, 57% of hand samples had Prevotellaceae,12% had Paraprevotellaceae, 5% had Succinivibrionaceae, and 2% at Spirochaetaceae (each at least 1% abundance). The majority of samples had some detectable reads from the genus Bifidobacterium, with 27% of samples comprised of at least 1% Bifidobacterium, which could be due to interaction with infant stool.

**Figure 2:**
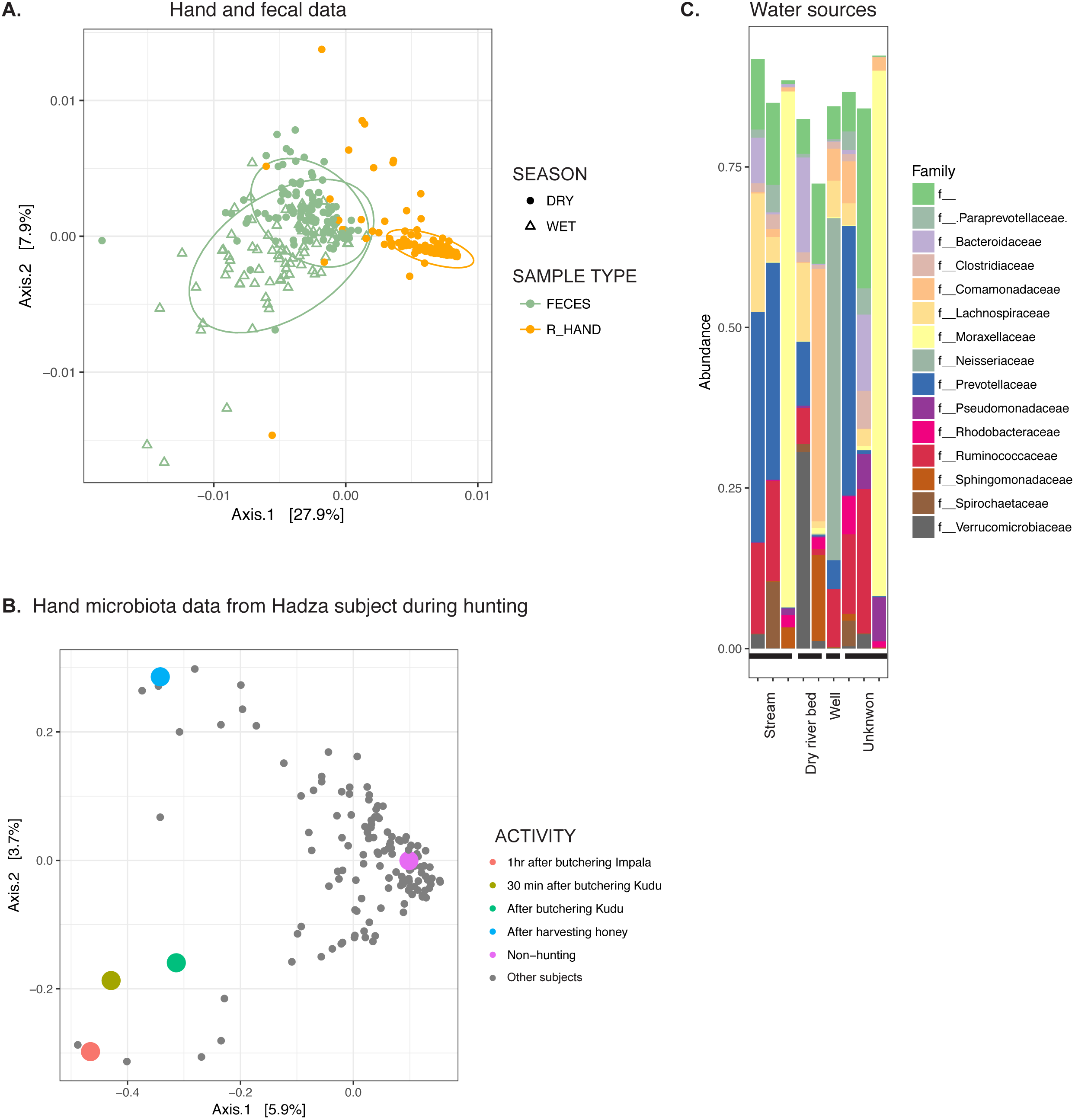
Microbes associated with hands, animals, and water sources in the Hadza environment. A. PCoA of weighted Unifrac distances of hand samples and fecal samples from Hadza. Fecal samples, green; hand samples, orange; closed circles, dry season; open triangles, wet season. Ellipses show .95 confidence level.
B. PCoA of unweighted Unifrac distances of hand samples. Colored dots represent samples acquired from the same subject; grey dots from remaining individuals. Colors indicate activities engaged in when sample was taken.
C. Composition of microbiota from water sources, summarized at the family taxonomic level. Families shown are limited to those present at greater than 1% in sum total of water samples.

Using information about daily activities corresponding to a subset of the samples, we addressed whether aspects of lifestyle could drive shifts in the hand microbial composition. Figure 2B shows a PCoA plot of unweighted Unifrac distances between hand samples from the entire cohort; one individual sampled at several different points is shown in color. On a non-hunting day, this individual’s hand microbiota clustered with the majority of the hand samples from the cohort. However, on two separate hunting days, one in which he butchered a lesser kudu *(Tragelaphus imberbis)*, another in which he butchered an impala *(Aepyceros melampus)*, his hand microbiota is an outlier to the group (Figure 2B). It should be noted that we do not have samples on hunting days prior to the kills, and therefore cannot be certain that this effect was due to contact with the animal versus other aspects of hunting days. Notably, the hand sample from a day he harvested honey (from honey bee *Apis mellifera scutelata)* is also an outlier, yet distinct from the hunting days (Figure 2B, blue circle). These data indicate each touchpoint on a forager’s landscape may affect the hand microbiota.

The data are consistent with the Hadza gaining exposure to distinct subsets of microbes via hunting and foraging. Although we are not statistically powered to answer the question directly of whether this exposure in the environment contributes to individuals’ microbial ecosystems, we analyzed honey samples taken from bee hives and stool, fur, and stomach swabs from animals in the environment. We were interested in determining whether these sources contained microbes found in the Hadza gut. To this end we inferred sequence variants from 16S amplicon sequencing data using the DADA2 method to enable matching exact sequences (amplicon sequence variants, or ASVs), rather than comparing OTUs, each of which contains a range of sequence variants(*19)*. We identified the ASVs shared in Hadza gut samples during the dry season (restricting our analyses to ASVs present in at least 10% of samples to improve confidence) that were taxonomically assigned to VANISH families and looked for them in the animal and honey samples. Interestingly, we found that of the shared Hadza gut ASVs from the four bacterial families, the majority are also present in at least one animal sampled (23/37 Prevotellaceae ASVs, 4/5 Spirochaetaceae ASVs, 5/8 Paraprevotellaceae ASVs, 2/3 Succinivibrionaceae ASVs). Animals sampled include dik dik (Madoqua sp.), lesser kudu (Tragelaphus imberbis), impala (Aepyceros sp.), hyrax (Heterohyrax brucei), zebra(Equus sp.), cow (Bos tauras), and vervet monkey (Chlorocebus pygerythrus); these ASVs were most often identified in the animal fecal samples. A similarly high rate ASV occurrence was observed in the bee hive samples (29/37 Prevotellaceae ASVs, 4/5 Spirochaetaceae ASVs, 6/8 Paraprevotellaceae ASVs, 2/3 Succinivibrionaceae ASVs). While not conclusive, these results support that interaction with animals and bee hives provides an environmental reservoir of the VANISH microbial families, a hypothesis that warrants detailed follow-up.

With a sanitized water supply serving as a major characteristic of industrialized society, we wondered whether the water sources available to the Hadza and other animals on the landscape may serve as an additional source of gut-colonizing microbes. We sampled water during the dry season from the surrounding area of the Hadza camps including streams, a well, and the dry riverbed. The samples varied in composition, but several had high levels of Prevotellaceae, one sample was 10% Spirochaetaceae, and several had low levels of Paraprevotellaceae (Figure 2C). When examining the same shared ASVs from the Hadza gut from the four VANISH families, nearly all were found in at least one water source (32/37 Prevotellaceae ASVs, 5/5 Spirochaetaceae ASVs, 7/8 Paraprevotellaceae ASVs, 2/3 Succinivibrionaceae ASVs). To address the possibility of contamination of water samples during sample preparation, we used an updated version of SourceTracker(*20*) to compare the probabilistic contribution of fecal samples proximal into water samples on the plate, relative to the contribution of randomly sampled fecal samples located on separate plates, and did not see any difference (p > 0.05, Wilcoxon test). While we are not equipped with sufficient data nor study design to explore the question of transmission, we found that Hadza gut bacteria that annually become undetectable and then re-appear are also present in the surrounding environment, making it a possible that the Hadza gut is repopulated via environmental sources.

## Hadza microbial diversity increases with age

We wished to investigate the development of the Hadza microbiota in the context of industrialized and other traditional populations, an important topic in light of known variation in human microbiota development*(21*). We analyzed the fecal samples from the adult Hadza(*1*) with additional samples obtained from Hadza children. When the OTUs of these samples are plotted together, we observed an increase in microbial diversity as the Hadza age (Figure 3A), particularly during the first few years of life, as has been reported in other traditional and industrialized cohorts(*11*). Interestingly, we do not see decreasing levels of diversity in the elderly Hadza as has been reported in some industrialized cohorts(*22*), although our sample size of elderly individuals is small. This difference may be due to the fact that the elderly Hadza continue to live in close proximity with the rest of the camp, which may help retain access to food, activities, and microbes characteristic of younger individuals. In industrialized cohorts the age-associated decline in diversity was most pronounced for elderly living in institutions, who are more segregated from free-living younger individuals. In contrast, a large study of healthy Chinese individuals showed little difference between individuals from age 30 to 100, and no decrease in alpha diversity(*23*). Future studies that investigate the relationships between diet, lifestyle and immune status are needed to determine whether specific microbial profiles are linked with healthy aging.

**Figure 3:**
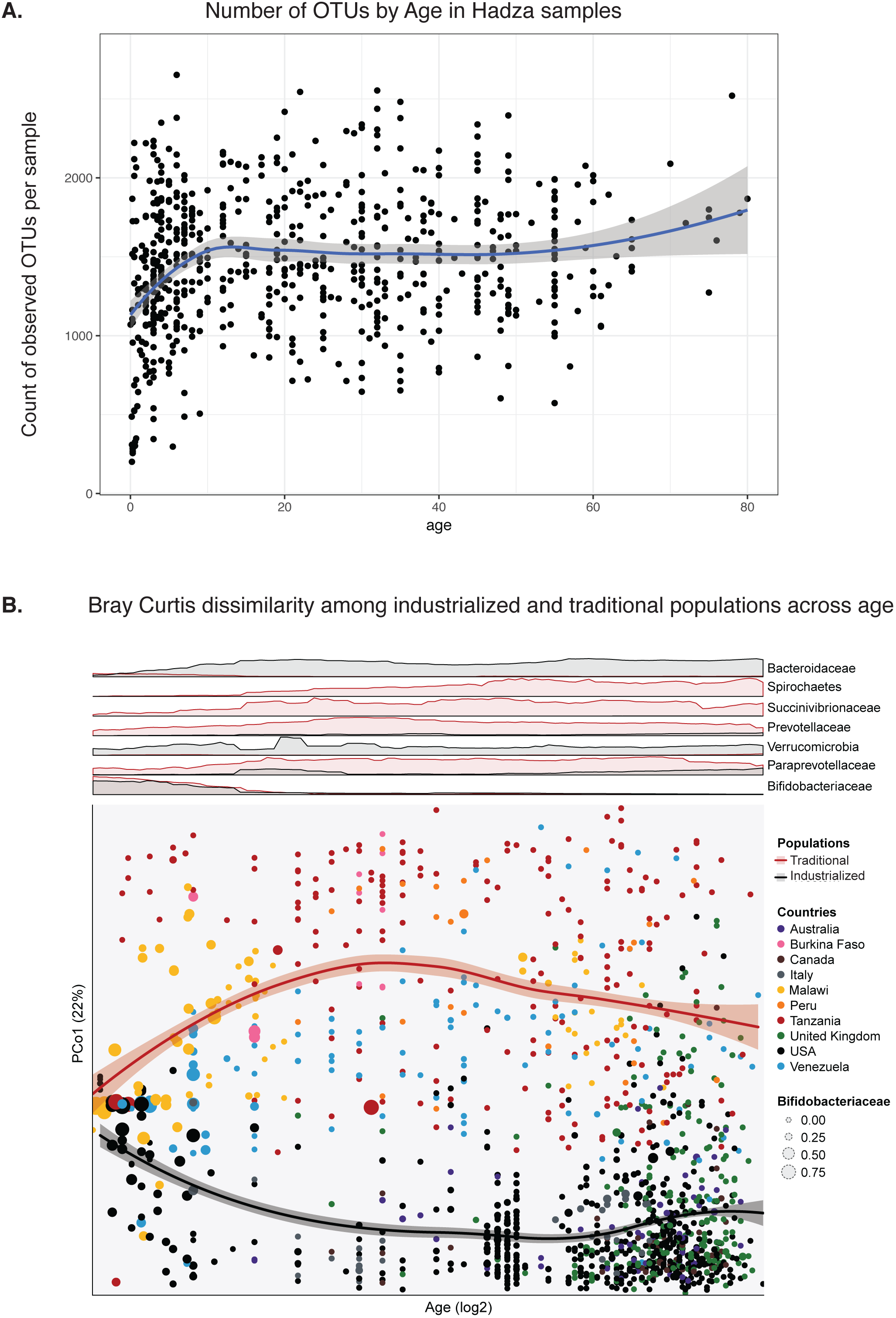
Hadza microbial diversity increases with age and diverges from industrialized populations. A. Number of OTUs detected in rarefied samples plotted by age. Trendline showing loess non-parametric regression line, standard error in grey.
B. Bottom: A scatterplot with axes Bray-Curtis dissimilarity principal coordinate 1 and Log2-transformed Age *(log2(Age + 1))* of microbial community compositions described at the family taxonomic level. The circles are colored by the country from which the subjects were sampled, and diameters based on relative abundance of Bifidobaceriaceae within the sample. Loess regression was applied to samples from industrialized and traditional populations using PCo1 coordinates and Log2-transformed Age with curves plotted according to the populations with 95% pointwise confidence interval bands. Top: Overlapping density plots (industrialized in black, traditional in red) representing the moving average of the relative abundance of families within the respective samples along the Log2-transformed Age x-axis and min-max scaled across both populations to allow for direct comparison.

We compared Hadza samples to data from other traditional cohorts and industrialized cohorts during the first few years of life (Figure 3B). Plotting Bray-Curtis dissimilarity, a metric of shared species between samples, we observe a high degree of similarity between populations early in life; however, cohorts with different lifestyles diverge with increasing age (Figure 3B, bottom panel). This divergence may reflect any number of differences between traditional and industrialized societies including higher consumption of complex carbohydrates, lower antibiotic use, and environmental exposure to a more diverse set of microbes in traditional populations relative to industrialized populations. Interestingly, the Hadza diverge almost immediately, and earlier relative to the other traditional and industrialized populations. Bifidobacteriaceae, a family commonly associated with the gut of breast-fed babies, occurs at high abundance and prevalence early in life then declines with age in both the traditional and industrialized cohorts (Figure 3B). In traditional populations, the VANISH families show a similar pattern of having low prevalence early in life and then increasing with age (Figure 3B, top panel).

## Limited sex differences in the Hadza microbiota

We explored differences between samples taken from male and female Hadza, since sexual division of labor characterizes the central-based foraging Hadza. Previous studies have described differences in diet between men and women(*24*), which is reflected in different dental wear patterns(*25*). We therefore wondered if sexual division of labor and potential macronutrient differences in diet documented among the Hadza resulted in differences in microbiota composition. Applying PCoA of weighted Unifrac distance did not statistically distinguish between male and female samples when using more Hadza samples than previous reports (Figure 4A)(*3*). Interestingly, there was no difference the data was subdivided by camp, despite different degrees of acculturation and sexual division of labor across camps. Consistent with the previous study, we observed a difference in the abundance of the genus Treponema between sexes, but not the genus Blautia (p = 0.031, p = 0.13, respectively, Wilcoxon test). However, when applying an unbiased approach using a statistical method for relative abundance data with multiple-hypothesis correction(*26*), the only genus significantly different was Dialister (Figure 4C, FDR < 5%; p = 0.0059, post-hoc Wilcoxon test). When restricting the analysis to only samples taken during the early wet season, we found the only significantly different genus to be Faecalibacterium (Figure 4D, FDR < 5%; p = 0.017, post-hoc Wilcoxon test). Therefore the characterization of the Hadza gut microbiota substantially differing by sex may need to be reconsidered.

**Figure 4:**
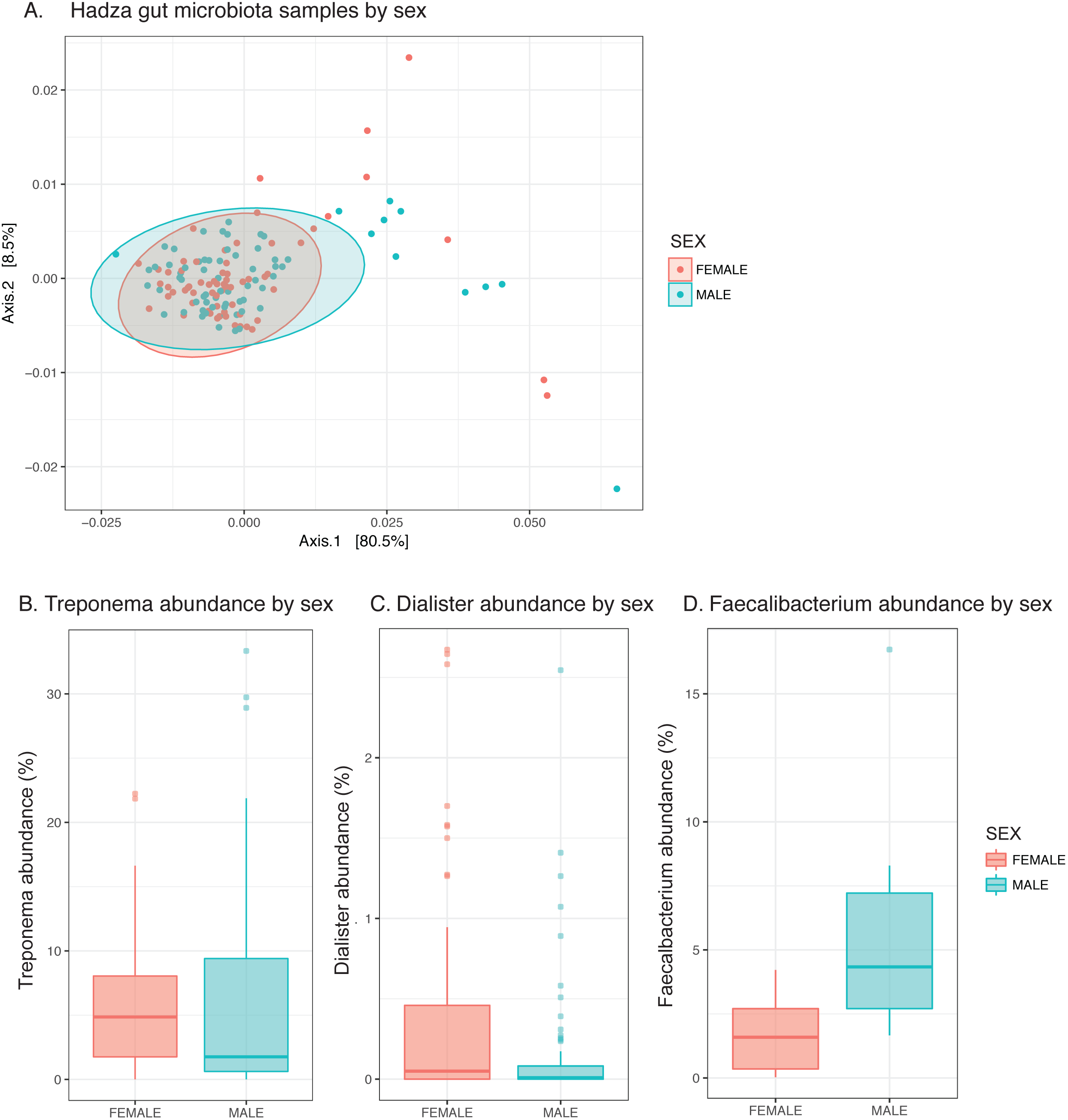
Limited sex differences in the Hadza microbiota. A. MDS plot of weighted Unifrac distance of Hadza fecal samples; pink, female; blue, male. Populations not significantly different (p-value = 0.328, permuted F-statistic).
B. Genus Treponema abundance by sex; p-value = 0.031, Wilcoxon test.
C. Genus Dialister abundance by sex; significant with FDR < 5% across all genera with ANCOM test; p-value 0.0059, Wilcoxon test.
D. Genus Faecalibacterium abundance by sex from samples taken from the early wet season to match sampling season from previous report; significant with FDR < 5% across all genera with ANCOM test; p-value 0.017, Wilcoxon test.

## The introduction of maize into the Hadza diet

Dietary perturbation has been shown to substantially influence the human microbiota(*27–29*). The Hadza consume a diet that is primarily composed of tubers, baobab, berries, honey, and hunted meat. This diet is distinct from the typical diet of industrialized societies. While it is difficult to isolate the effect of diet versus other lifestyle and geographical differences that distinguish the Hadza from other populations, we were able to sample individuals that underwent a temporary but significant shift in diet. In the Ukamako camp, individuals were consuming primarily baobab, roots, berries, and honey until they received a large bag of unmilled maize on January 30^th^, 2014 (“Day 0”). During the days following, maize was consumed for breakfast, lunch, and dinner. We analyzed fecal samples in the days leading up to and following the maize arrival for twelve individuals; eight of these individuals were sampled in all five sequential days prior to and during maize consumption.

Plotting the first two principal components of a principal component analysis of unweighted Unifrac distances revealed a shift in the first principal component in the 24hrs and 48hrs post-maize consumption, relative to the days pre-maize consumption in most individuals (Figure 5A, 5B). When we quantified unweighted Unifrac distances between each time interval within the eight individuals sampled for five consecutive days, the microbiota perturbation induced by the maize diet became apparent. Day 0 and Day 1 (pre-maize and post-maize) samples exhibited a larger distance compared to Day −2 and Day 0 (two pre-maize time points) (Figure 5C, p = 0.0078, Wilcoxon paired test).

**Figure 5:**
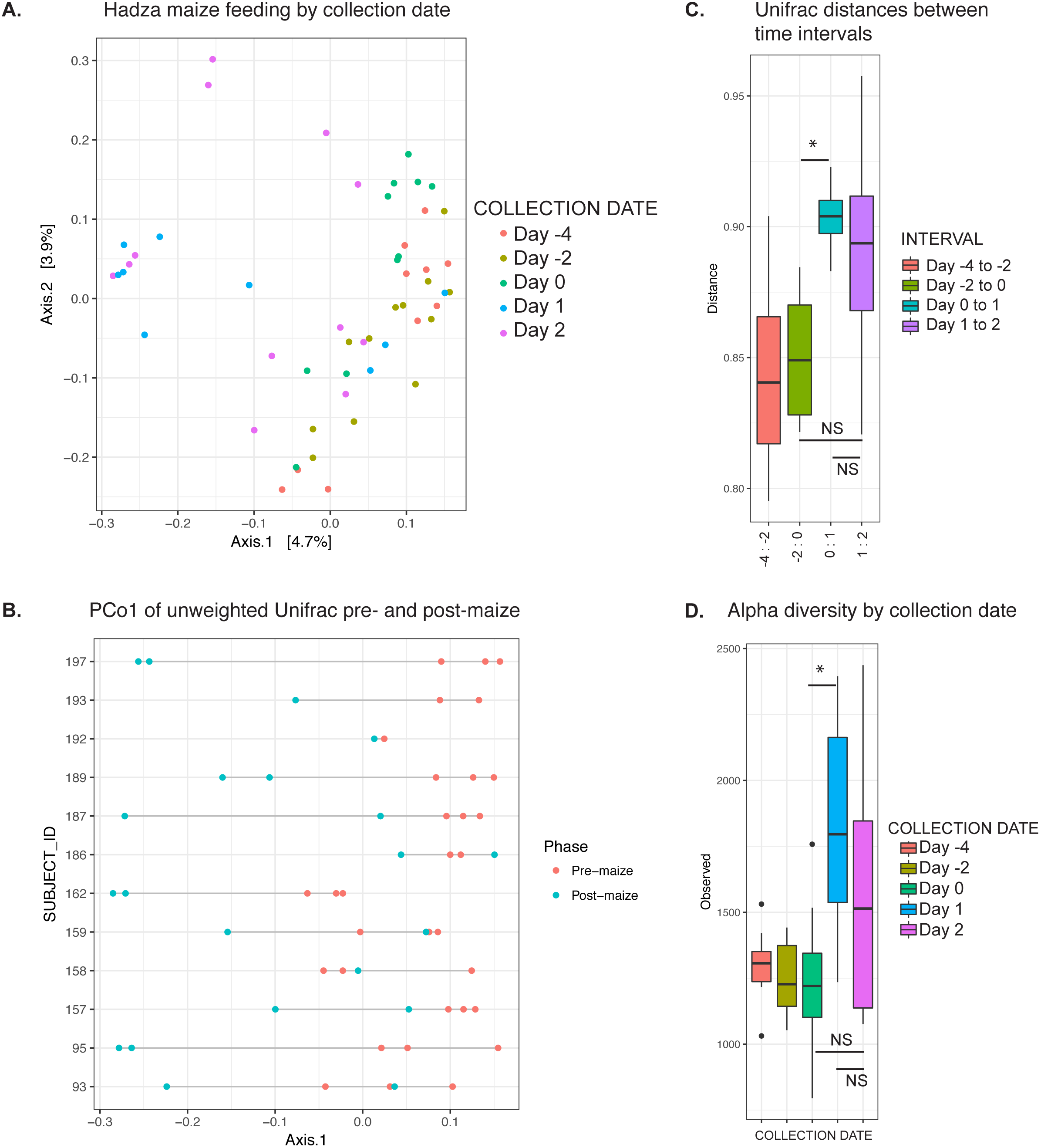
The introduction of maize into the Hadza diet. A. PCoA of unweighted Unifrac distances of fecal samples taken during maize introduction. Samples labeled by collection date. Maize introduction on Day 0, 1/31/14. Day −4, 1/26/14; Day −2, 1/28/14; Day 1, 1/31/14; Day 2, 2/1/14.
B. PC1 from PCoA of unweighted Unifrac distance plotted by subject. Points per sampling time. Teal dots are pre-maize (Day −4, −2, 0), red dots are post-maize (Day 1, Day 2).
C. Unifrac distance between time intervals. * p-value = 0.0078, Wilcoxon paired test.
D. Observed OTUs per rarefied sample, grouped by collection date. * p-value = 0.016, Wilcoxon paired test.

An analysis of alpha diversity revealed an increase in observed OTUs 24hrs after the maize arrival (Day 0 vs. Day 1, p = 0.016, Wilcoxon paired test), while the second day was not significantly different from the two preceding days (Figure 5D). These data are consistent with observations from a short-term controlled feeding experiment that found a greater perturbation in the first 24 hours than in the days following(*28*). While more sampling is needed to address potential longer-term changes to the microbiota, these data suggest a change in diet produces rapid changes in the microbiota in the Hadza.

## Conclusion

The gut microbiota has emerged as a critical modulator and indicator of human health and disease. While many associative and causal links have been established between the microbiota and health outcomes, the microbial taxa, functionality, or metabolic states that are either protective or drive the development of disease are poorly understood. Looking to populations across the world that have been minimally affected by industrialization can serve as a model for identifying critical organisms and functionality that have been lost in industrialized populations. Through an analysis of the microbiota from the Hadza hunter-gatherers and integrating their data with other populations, Smits et. al identified bacterial families that have been maintained in traditional societies across the world but are diminished or lost with modernization. These bacterial taxa are candidates for future study toward a better understanding of the co-speciation of humans and gut microbes, and of what has been disrupted in recent times.

Despite seasonal fluctuations of bacterial taxa, the Hadza are able to maintain a diverse microbiota over sequential years with species returning during the dry season that were undetectable in the wet season. These microbes may be present in the gut below our level of detection or they may be absent and then reintroduced to the gut, perhaps from reservoirs in the environment. In working toward a better understanding of the regional species pool, this study serves as a first survey of the Hadza environmental microbiota and the concordance with the organisms appearing in the gut. Though the analysis is limited by sample number and interval to make broad claims, it provides case studies of how the environment may impact the cycle of the Hadza microbiota. Forthcoming studies will provide more granularity by season, activity, and environmental site and how these correspond to the Hadza-resident microbiota.

In addition to the influence of the seasons and the environment, we have observed how the microbiota changes with age and how it compares across sex. We observed short-term changes associated with the introduction of maize into the diet, an important area to examine as industrialization brings with it dramatic changes in diet. We hope this work serves a snapshot of the state of the Hadza microbiota in the context of environment, diet, and lifestyle that can inform our understanding of the microbiota across a diverse set of populations.

## Methods

Data was generated and analyzed as previously described(*1*). Diversity analyses and ordination was performed using the R package phyloseq. DADA2 analysis was performed using the R package dada2. An updated form of SourceTracker https://github.com/biota/sourcetracker2 was used to examine contamination in sample preparation. Comparison was performed between the contribution to the water samples from the eight closest wells on the plate, relative to eight fecal samples sampled randomly from the remaining plate locations. Distance comparison between sexes was performed using the adonis function in the R package vegan. Genera comparisons between sexes were done using the R package ancom.R.

